# Super-Resolution Diffusivity Mapping Reveals Spatial Correlations Between Lateral Mobility and Structural Heterogeneities on Cellular Membranes

**DOI:** 10.1101/2025.04.01.646704

**Authors:** Chu Han, Zhe Zhao, Huihui Gao, Jingwen Deng, Qi Wang, Yihan Wang, Liu Liu, Lin Xu, Shuoxing Jiang, Xiang Zhan, Kun Chen, Rui Yan, Wan Li, Weihua Huang, Ke Xu, Limin Xiang

**Affiliations:** College of Chemistry and Molecular Sciences & Taikang Center for Life and Medical Sciences, Wuhan University, Wuhan, Hubei 430072, China; State Key Laboratory of Coordination Chemistry, Department of Biomedical Engineering, College of Engineering and Applied Sciences, Nanjing University, Nanjing, Jiangsu 210023, P. R. China; Department of Biostatistics, Peking University, Beijing 100871, China; School of Optoelectronic Science and Engineering, University of Electronic Science and Technology of China, Chengdu, Sichuan 610054, China; Department of Genetics, Blavatnik Institute, Harvard Medical School, Boston, MA 02115, USA; Department of Chemistry, University of California, Berkeley, CA 94720, USA; Department of Physics, University of Muenster, Muenster, 48159, Germany

## Abstract

Cellular membranes orchestrate critical processes such as molecular transport and signal transduction, both regulated by the lateral mobility of lipids and proteins. However, resolving nanoscale diffusional heterogeneities and elucidating their underlying mechanisms remains a formidable challenge due to the membrane’s intricate architecture and compositional diversity. Here, we present point-cloud single-molecule diffusivity mapping (pc-SM*d*M), a cutting-edge super-resolution technique that offers a point-cloud data format with enhanced spatial resolution for diffusivity mapping. Using pc-SM*d*M, we visualize nanoscale diffusion slowdown clusters with ∼50 nm in diameter on plasma membranes. These clusters are predominantly governed by cholesterol content, alongside contributions from membrane protein assemblies and topographical features. Leveraging two-color pc-SM*d*M, we concurrently imaged multiple lipids and membrane probes, revealing distinct "fingerprint" diffusivity maps shaped by their interactions within lipid bilayers. Our findings position pc-SM*d*M as a transformative tool for spatially resolving molecular interactions and membrane dynamics in live cells, offering new insights into the underlying mechanisms that govern membrane mobility at the nanoscale.

## Main

Biological membrane is a crucial lipid bilayer that enables many critical cellular processes such as signal transduction^1, 2^ and molecular transport^3^. The mobility of the lipid bilayer highlights the importance of quantifying biomolecular diffusion on the membranes, as diffusion rates fundamentally limit these processes^4^. However, deciphering the molecular basis of diffusional behavior on the plasma membrane poses significant challenges due to its complex nanoscale architecture, characterized by its dynamic structural variations and heterogeneity in composition. Lipid rafts^5–7^, compact membrane domains rich in saturated lipids and cholesterol and typically spanning 50-200 nm, feature prominently in determining membrane mobility^8, 9^. Nanoscale assemblies of membrane proteins, ranging from tens to hundreds of nanometers in size, have been identified as impediments to molecular diffusion on the plasma membrane^10, 11^. Furthermore, the presence of blebs or bleb-like protrusions adds a third dimension to consider, potentially compartmentalizing lateral diffusion on the two-dimensional lipid bilayer^4, 12^. Consequently, a detailed understanding of diffusion behaviors, accounting for these sub-diffraction-limited (<∼300 nm) compositional and topographical variances with high spatial resolution is essential.

Lateral mobility on plasma membranes has been studied using various diffusion measurement techniques including FRAP (fluorescence recovery after photobleaching)^13^, FCS (fluorescence correlation spectroscopy)^14, 15^, and SMT (single-molecule tracking)^16–19^. The most elementary mode of diffusion on a perfectly homogeneous membrane is Brownian motion, characterized by a mean squared displacement proportional to time. However, experimental evidence has revealed that various obstacles disrupt Brownian motion on the plasma membrane, giving rise to complex diffusional behaviors such as transient anchorage, confined diffusion, and hopping. The ’Anchored Picket Fence Model’ is a prominent framework that describes this complexity^2^. This model proposes that the cytoskeleton forms a ’fence’ beneath the plasma membrane, creating compartments, while transmembrane proteins serve as ’pickets’. Kusumi and colleagues demonstrated this concept through SMT studies tracking single lipid movements on the plasma membrane. While their use of large probe particles has sparked some debate^20^, subsequent studies employing smaller fluorescent lipids has partially validated their model^21^, but suggesting a milder confinement effect by the ’fence’. Despite these insights, a direct spatial correlation between local diffusional confinements and the corresponding ultrastructures on cellular membranes is still needed to fully elucidate the mechanisms underlying heterogeneous diffusion.

Recent advances in sub-diffraction-limited optical imaging have significantly enhanced our capability to visualize nanoscale membrane ultrastructures^22–24^. Among these, single-molecule localization microscopy (SMLM) has emerged as a powerful technique, enabling super-resolved localizations of individual molecules to generate high-resolution, point-cloud images from millions of events within a wide-field view^25–28^. Beyond spatial localization, SMLM leverages additional molecular properties, such as spectra and displacements, to construct multidimensional images enriched with physicochemical information in live cells^29^. Unlike conventional pixelated images, point-cloud representations offer enhanced flexibility in image processing and data analysis^30^, allowing precise drift correction, rotation, and seamless correlation across all SMLM datasets^31^. Yet, the absence of a nanoscale diffusivity mapping method compatible with point-cloud formats has limited the exploration of diffusional dynamics on cellular membranes. Techniques such as STED (stimulated emission depletion)-FCS achieve high spatiotemporal resolutions^32^, but are constrained to pixelated and one-dimensional mapping^33^. Similarly, single-molecule tracking (SMT) excels in elucidating diffusional behaviors but suffers from low throughput, making it unsuitable for spatial diffusivity mapping^34^. While our single-molecule diffusivity mapping^35^ (SM*d*M) achieved significant strides in pixelated representations of diffusivity (∼100 nm bin size), it falls short in achieving the spatial resolution and point-cloud fidelity necessary to unravel finer details and the underlying mechanisms driving diffusion heterogeneities.

To overcome these limitations, we introduce point-cloud single-molecule diffusivity mapping (pc-SM*d*M), an advanced technique that applies a point-cloud-specific Gaussian filter^36^, preserving data integrity while enhancing spatial resolution and image quality. This approach facilitates direct visualization of nanoscale diffusion slowdown clusters (∼50 nm in diameter) on plasma membranes and enables correlative analysis with all other SMLM datasets, shedding light on molecular interactions involving cholesterol, membrane proteins, and topographical geometries. Furthermore, through concurrent diffusivity imaging of multiple lipids, pc-SM*d*M reveals distinct diffusional behaviors influenced by their interactions with lipid bilayers. These findings underscore pc-SM*d*M’s transformative potential to advance our understanding of molecular interactions and membrane dynamics at the nanoscale in live cells.

## Results

### pc-SM*d*M Imaging of Membrane Probes on Live Cell Membranes

For diffusivity mapping on cellular membranes, we employed the commercial dye BDP TMR azide, known for its strong affinity for lipid bilayers and superior performance in single-molecule imaging^37^. We introduced ∼3 nM of BDP TMR azide into the live cell medium prior to imaging. Like other hydrophobic membrane dyes, BDP TMR azide demonstrates an approximate tenfold increase in brightness when in hydrophobic environments as opposed to aqueous ones, rendering it highly effective for PAINT-type super-resolution imaging^38^. Its transient binding to the membranes yields strong fluorescent signals, enabling single-molecule tracking over multiple frames (Fig. 1a). The eventual photobleaching or unbinding of the dye reduces its brightness, thereby facilitating a proper control of molecular density necessary for single-molecule localization. To carry out single-molecule diffusivity mapping (SM*d*M), we excited the dyes with stroboscopic pulses of τ = 2 ms. By illuminating at the middle of each frame and a slightly smaller imaging area, we shorten the frame-to-frame separation time to ∼6 ms, thus allowing for the accumulation of single-molecule displacements between all consecutive frames (Fig. 1a). Through the acquisition of 65,000 frames, we identified approximately 2-4 million of molecules, recording a million of single-molecule displacements in total.

**Fig. 1:**
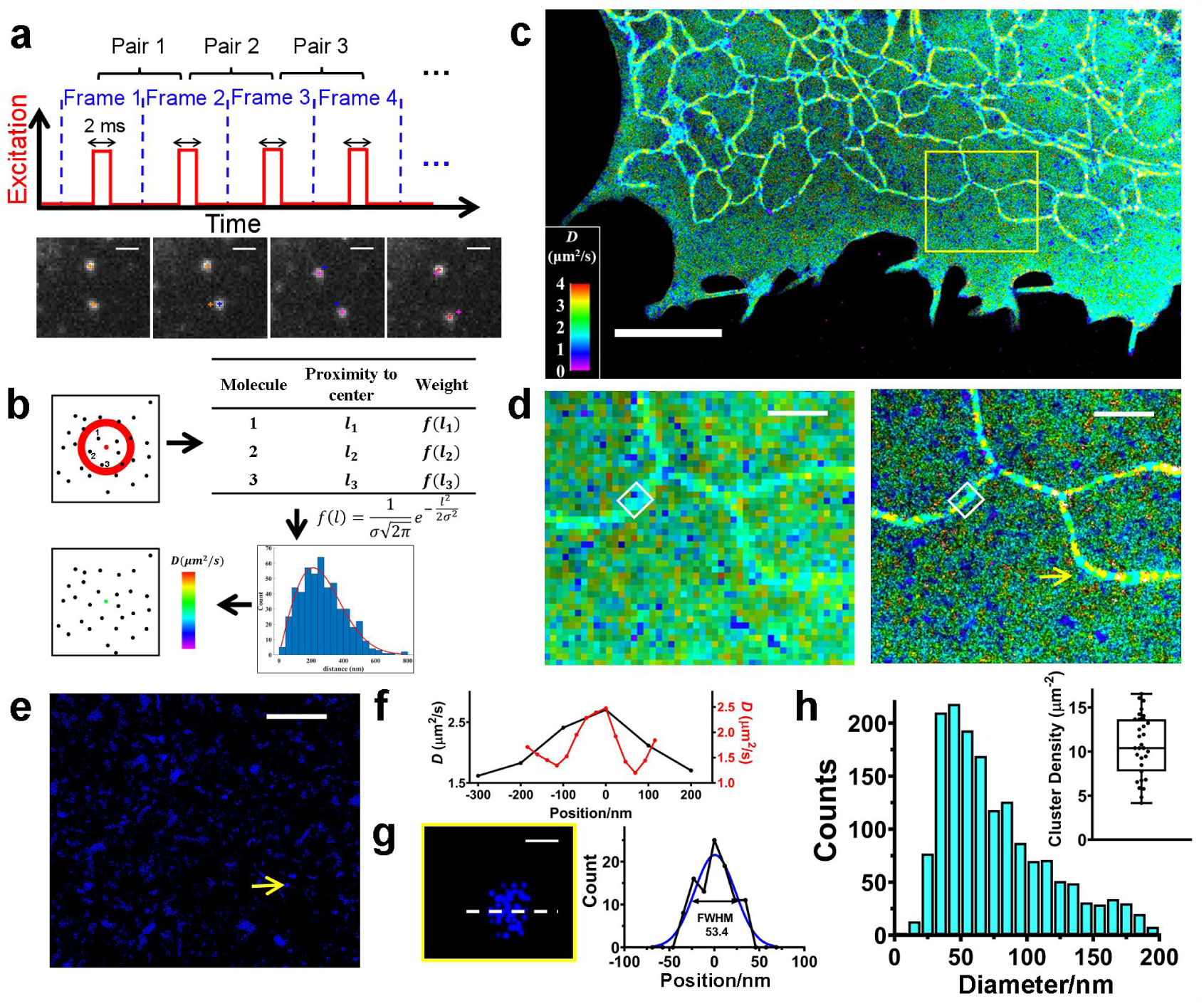
Super resolution mapping of diffusivity on cellular membrane via point-cloud single-molecule diffusivity mapping (pc-SM*d*M). **a,** Stroboscopic illumination with excitation pulses τ = 2 ms and frame-to-frame separation time of ∼6 ms, enabling efficient tracking of the membrane probe BDP-TMR-azide on the cellular membrane across multiple frames. **b,** In pc-SM*d*M, diffusion rates (*D*) were calculated based on the displacements (*d*) of surrounding molecules and assigned to the central molecule, preserving the point-cloud image format (see text for details). **c,** Point-cloud SM*d*M image of a live COS-7 cell labeled with BDP-TMR-azide. **d,** Comparison of pixelated (left) and point-cloud (right) SM*d*M images, highlighting the zoomed-in region indicated by the yellow box in panel **c**. Pixel size: 100 nm. **e,** Extraction of clusters with slow diffusion (slow-*D* clusters) from the images in panel **d**. **f,** Diffusion rate profiles plotted as a function of distance for an ER tubule, comparing pixelated (black) and point-cloud (red) SM*d*M images. Bin size for point-cloud data: 20 nm. **g,** Gaussian fitting of a representative slow-*D* cluster (indicated by yellow arrows in panels **d** and **e**) yields a full width at half maximum (FWHM) of 53.4 nm, reflecting the cluster diameter. **h,** Size distribution and density of slow-*D* clusters on the plasma membrane, derived from analyses across 20 regions in 10 cells. Scale bar: 1 μm in **a**, **d** and **e**, 5 μm in **c**, 100 nm in **g**. The color bar for diffusion rates is consistent across all panels. Box plot elements for all the figures: center line, median; box limits, upper (75%) and lower (25%) quartiles; points, outliers.

Next, we produced spatially-resolved diffusivity images by extracting diffusion rates from the displacements within designated regions. In our previous SM*d*M approach, accumulated displacements (denoted as "*d* values") were organized into spatial bins, typically spanning 100×100 nm^2^, to derive local diffusion rates (referred to as "*D* values") for each bin, resulting in a pixelated diffusivity mapping. In contrast, our current approach adopted a more refined strategy. Instead of grouping displacements into predefined bins, we examine the displacements of molecules surrounding each target molecule, usually within a radius of ∼50 nm. These displacements were utilized to construct histograms, but incorporating weighted values, calculated by the distances of neighboring molecules from the central molecule (*f* (*l* ) in Fig. 1b). Subsequently, the histogram was constructed based on these weighted values, and the derived diffusion rate (*D* value) was assigned to the central molecule, thus preserving the point-cloud nature of the super-resolved image (Fig. 1b).

Fig. 1c illustrates the diffusivity mapping of BDP TMR azide in a live COS-7 cell, showcasing the point-cloud SM*d*M (pc-SM*d*M) image. For endoplasmic reticulum (ER) tubule, we conducted an analysis of displacements along the one-dimensional structure, employing principal component analysis (PCA). This allowed us to accurately capture the diffusion rate along the ER tubule as the *D* value^39^ (Extended Data Fig. 1b-f). Notably, diffusion rates on the plasma membrane (PM) were found to be slower than those along the ER tubule, attributed to the higher cholesterol content in the plasma membrane^39^. The pc-SM*d*M image, in contrast to the pixelated version, unveils finer details of both the plasma membrane and the ER tubule. The improved resolution is evident in Fig. 1d, which zooms into the area marked by the yellow box in Fig. 1c. Subtle diffusivity variations across the plasma membrane and ER are clearly resolved, with the distinct morphology of the ER tubules prominently captured. These observations are further corroborated by the diffusivity profile plotted against the distance across an ER tubule (Fig. 1f, corresponding to the white boxes in Fig. 1d), where the point-cloud format of the pc-SM*d*M image allows for an adjustable bin size, enabling more detailed and precise analysis.

In-depth analysis of molecule diffusion on the plasma membrane (PM) revealed numerous clusters where diffusion markedly decreases, as highlighted by the yellow arrows in Fig. 1d&e. These variations suggest alterations in the underlying membrane structures. To quantify these observations, we utilized density-based clustering (DBSCAN) to evaluate the prevalence of clusters with diffusion rates below 1.27 μm^2^/s across 10 different cells (Extended Data Fig. 1a). Molecules exhibiting *D*<1.27 μm^2^/s were aggregated to visualize all slow-*D* clusters (Fig. 1e), with the full width at half maximum (FWHM) of each cluster representing its diameter (Fig. 1g). Statistical analysis (Inset of Fig. 1h) revealed an average of 10.2 ± 3.2 slow-*D* clusters per μm^2^ on the PM, primarily with diameters around 52 nm (Fig. 1h). Thus, with enhanced spatial resolution, point-cloud Single-Molecule Diffusivity Mapping (pc-SM*d*M) technique enables us to delineate diffusivity heterogeneity on plasma membrane with high precision. Although this resolution is contingent upon local single-molecule counts and the extent of local diffusion variations, the current advancement facilitates the direct wide-field imaging of picket-like clusters that impede diffusion on the PM.

### Cholesterol’s Influence on Lateral Membrane Mobility

We further investigated diffusivity on the nuclear envelope (NE), which shares a similar lipid composition with the ER^40^. Fig. 2a presents pc-SM*d*M image of the NE’s ventral side, offering exceptional spatial resolution that enables the visualization of nuclear pores for live cells. Beyond traditional super-resolution imaging (white box in Fig. 2a), our method distinctly captures the variation in diffusion rates between the NE and the nuclear ER. This difference is further detailed in the enlarged section of Fig. 2b (corresponding to yellow box in Fig. 2a), showcasing marginally slower diffusion on the nuclear ER compared to the NE. Additionally, we observed sporadic clusters with diffusion slowdowns on the nuclear ER (white arrow in Fig. 2b). Drawing from our prior findings, in which we attributed slower diffusion rates on the peripheral ER to ER-PM contact sites^39^, here we speculated that the observed diffusion deceleration on the nuclear ER may be associated with the nuclear envelope-ER juncture, a topic we will explore in greater depth subsequently.

**Fig. 2:**
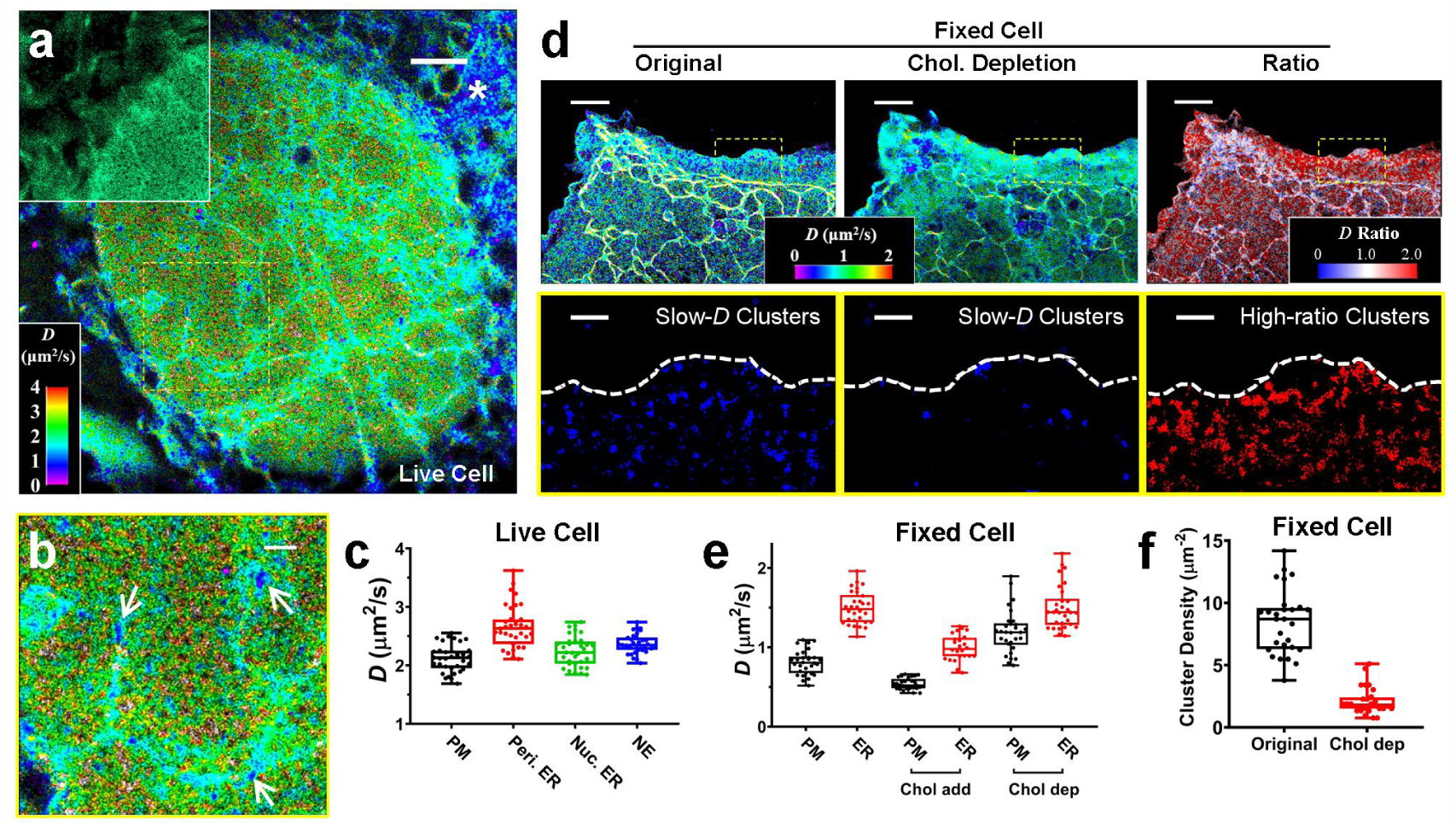
Cholesterol predominantly regulates the diffusivity on cellular membranes. **a,** pc-SM*d*M using BDP-TMR-azide depicting diffusion rates on biological membranes near the nucleus. Diffusivity is highly heterogeneous, with slower diffusion observed on nuclear ER membranes compared to the nuclear envelope. Inset: SMLM image of the nuclear membrane. **b,** Zoomed-in image of the yellow boxed region in panel **a**, showing finer ultrastructures including nuclear pores and nuclear ER. **c,** Statistical comparison of diffusion rates across various biological membranes, derived from analyses across 30 regions in 10 cells. **d,** Diffusivity mapping for fixed cells under baseline conditions (left), after cholesterol depletion (middle), and the corresponding ratiometric map (right). Cholesterol depletion significantly increases diffusion rates on the PM but not on the ER. **e,** Statistical analysis of diffusion rates on ER and PM under conditions of cholesterol addition and depletion in fixed cells, derived from analyses across 30 regions in 10 cells. **f,** Density analysis of slow-*D* clusters on the PM before and after cholesterol depletion in fixed cells, derived from analyses across 30 regions in 10 cells. Scale bar: 2 μm in **a** and upper panels in **d**, 500 nm in **b** and lower panels in **d**.

Fig. 2c summarized the typical diffusion rates across various cellular membranes, showing 2.7 ± 0.37 μm^2^/s and 2.4 ± 0.17 μm^2^/s for peripheral ER and the NE respectively, in contrast to 2.2 ± 0.26 μm^2^/s and 2.1 ± 0.23 μm^2^/s for the nuclear ER and PM respectively. Despite the nuclear and peripheral ER having comparable lipid compositions in their membranes, the diffusion rates on the nuclear ER membrane are subtly lower. This decrement is likely due to a higher density of ribosomes/proteins on the nuclear ER membrane, thereby creating a more crowded environment. In addition, the ER and mitochondria encircling the nucleus are densely packed, making accurate measurements of their diffusion rates challenging (asterisk in Fig. 2a).

Cholesterol is known to be rich in PM comparing to ER/NE, and it assists the organization of lipid bilayers into more ordered and tightly packed phases. To quantitatively assess the impact of local cholesterol concentrations on slow-*D* clusters within the PM, we performed pc-SM*d*M imaging before and after the depletion of cholesterol using methyl-*β*-cyclodextrin (M*β*CD)^41^. Due to the morphological changes live cells undergo during treatment, we fixed the cells prior to imaging, and the fixation caused a global slowdown in diffusion rates. Fig. 2d illustrates the diffusivity maps on plasma membranes before and after cholesterol depletion (left and middle panels), revealing that the treatment led to an approximate 40% increase in plasma membrane diffusion rates, but less impacts on the ER. This suggests a more pronounced impact of cholesterol depletion on plasma membranes. Conversely, adding cholesterol through cholesterol-M*β*CD complexes resulted in a greater reduction in diffusion rates on the ER compared to the plasma membrane, as shown in Fig. 2e and Extended Data Fig. 2a. Cholera Toxin Subunit B (CTB) treatment in a live cell also induced the formation of nanodomains with high contents of cholesterol on the plasma membrane^39^, leading to local diffusion slowdowns (Extended Data Fig. 2b). Our method’s point cloud feature enabled us to generate a ratiometric image by comparing diffusion rates before and after cholesterol treatment (see Methods for details). As illustrated in Fig. 2d (right panel), the ratiometric image reveals a markedly higher diffusion rate ratio on the plasma membrane (PM) compared to the endoplasmic reticulum (ER) following cholesterol depletion. A closer examination of selected areas (yellow box in Fig. 2d) reveals that cholesterol depletion eliminated some, but not all, slow-*D* clusters, indicating that factors beyond cholesterol may contribute to these slow-*D* clusters. Subsequent application of DBSCAN across 10 different cells helped quantify these effects, revealing a significant reduction in the prevalence of clusters with diffusion rates lower than 0.37 μm^2^/s. The comparative analysis, depicted in Fig. 2f, quantifies these clusters per μm^2^ on the PM of fixed cells before and after cholesterol depletion, with decreasing densities from 8.5 ± 2.6 per μm^2^ to 2.1 ± 1.1 per μm^2^, highlighting cholesterol’s pivotal role in modulating membrane mobility.

### Role of Membrane-Associated Proteins in Regulating Lateral Membrane Mobility

After identifying cholesterol as a predominant lipid component influencing diffusion, we next investigated the influence of membrane-associated proteins on the lateral mobility of the plasma membrane (PM). We performed pc-SM*d*M on live cells, followed by immunofluorescence labeling of the targeted proteins in fixed cells. Two anchoring proteins, ankyrin-B (ANK2) and adducin, were selected for this study, as both are involved in maintaining the structural integrity of the PM by anchoring membrane proteins to the cytoskeleton^42^. Fig. 3a and Fig. 3b show the pc-SM*d*M of cellular membrane and SMLM images of adducin, respectively. Cross-correlation analysis^43^ revealed a weak overall correlation between slow-*D* clusters and adducin across the entire image (blue curve, Fig. 3c). However, in specific regions (as shown in the zoom-in images of Fig. 3c), the correlation became notably stronger (red curve, Fig. 3c). Similar trends were observed for ankyrin-B, where the overall correlation remained weak, but certain areas exhibited stronger correlation (Extended Data Fig. 3a-c). These results suggest that the underlying membrane-anchored proteins impede diffusion on the plasma membrane in a few localized regions, aligning with previous findings from experiments involving small fluorescent probes^21^.

**Fig. 3:**
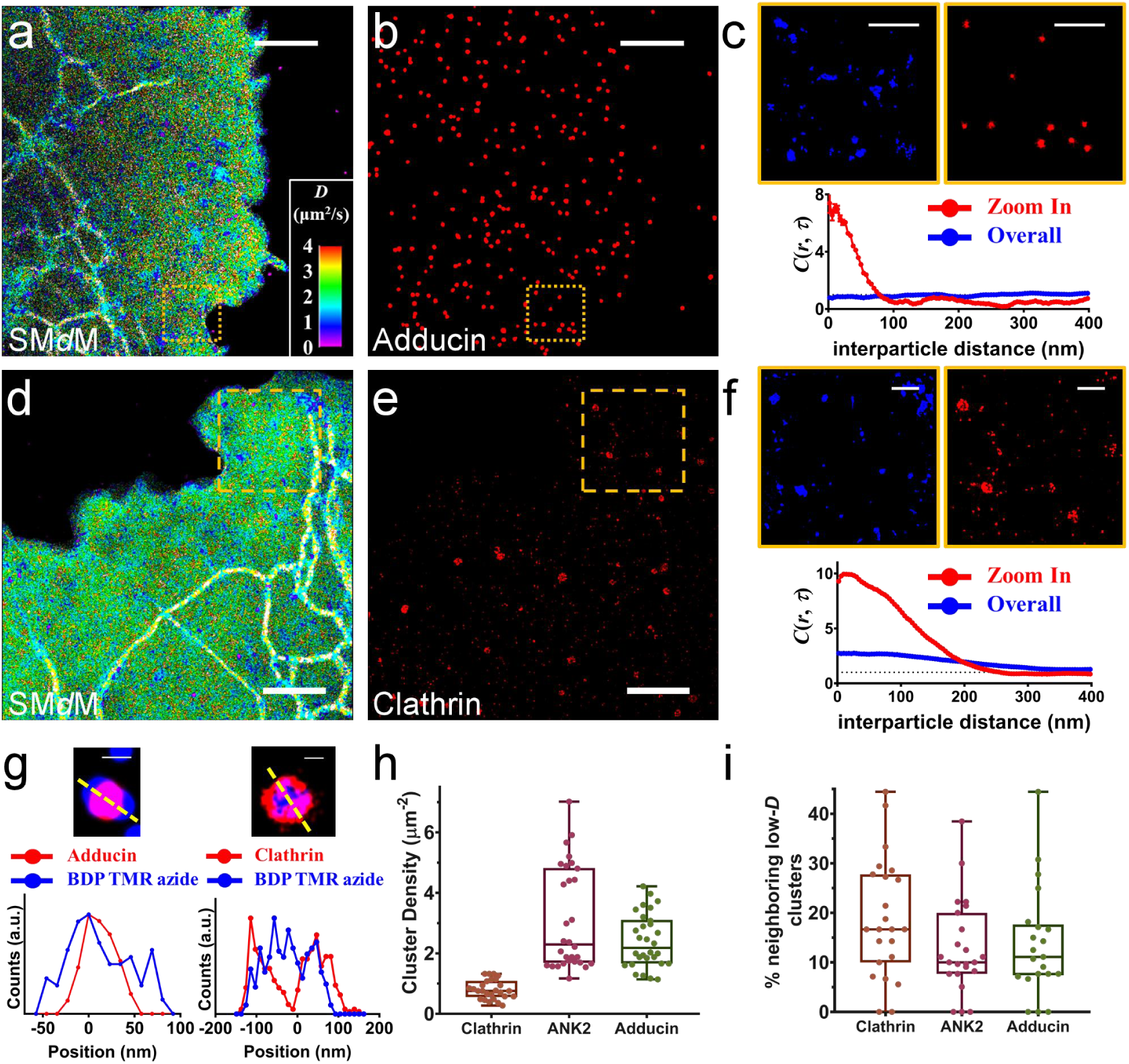
Membrane-associated proteins impede diffusion on the plasma membrane in some localized regions. **a-b,** Correlative pc-SM*d*M image of a live cell using BDP-TMR-azide and SMLM image of adducin. **c,** Enlarged views of the green boxed regions in panels **a** and **b**. Blue clusters indicate slow-*D* clusters on the PM. Cross-correlation analysis reveals strong correlation within the zoomed-in area (red curve) but not across the entire image (blue curve). **d-e,** Correlative pc-SM*d*M image of a live cell and SMLM image of clathrin. **f,** Enlarged views of the green boxed regions in panels **d** and **e**. Blue clusters indicate slow-*D* clusters on the PM. Similar to adducin, cross-correlation analysis shows good correlation within the zoomed-in area (red curve) but not across the entire image (blue curve). **g,** Cross-sectional profiles from merged images of adducin and clathrin with slow-*D* clusters, highlighting distinct correlative patterns. **h,** Cluster density of membrane-associated proteins clathrin, ankyrin-B, and adducin, derived from analyses across 30 regions in 10 cells. **i,** Likelihood of finding a neighboring slow-*D* cluster for clathrin, ankyrin-B, and adducin protein clusters, derived from analyses across 25-30 regions in 8-10 cells. Scale bar: 2 μm in **a**, **b**, **d**, and **e**, 500 nm in **c** and **f**, 100 nm in **g**. The color bar for diffusion rates is consistent across all panels.

Endocytic scaffolding proteins such as clathrin assist in the formation of lipid domains with distinct components and curvatures during endocytosis^44^. Recent studies have also suggested that clathrin-coated pits similarly impede protein diffusion^10, 45^. To determine whether endocytic scaffolding proteins reduce membrane diffusion for small molecules, we performed correlative imaging of diffusivity and clathrin (Fig. 3d,e). Despite the lower density of clathrin clusters on the PM, cross-correlation analysis between slow-*D* clusters and clathrin exhibited a relatively stronger correlation across both the entire image and specific areas, when compared to adducin (Fig. 3f, note the *y*-axis scale difference). A detailed comparison of the merged images for adducin and clathrin revealed different correlation patterns (Fig. 3g). Slow-*D* clusters were observed to typically surround adducin clusters, exhibiting a relatively larger diameter. In contrast, slow-*D* clusters were sometimes enclosed within hollow clathrin-coated pits. These findings suggest that these two proteins influence membrane diffusion through different mechanisms: while anchoring proteins like adducin slow local diffusion by ’nailing’ pickets into the membrane, endocytic scaffolding proteins generate confined lipid domains, potentially altering both lipid composition and 3D geometry. A quantitative comparison of these membrane-associated proteins is presented in Fig. 3h and Fig. 3i. Ankyrin-B and adducin exhibited higher cluster densities than clathrin. However, clathrin demonstrated a greater likelihood of encountering neighboring slow-*D* clusters compared to ankyrin-B and adducin, further underscoring its stronger influence in impeding membrane diffusivity. Similar patterns were observed between the diffusivity on fixed cell membranes and these protein nanodomains, effectively ruling out potential artifacts caused by cellular motion during fixation (Extended Data Fig. 3d-i).

### Three-Dimensional Topographical Effects on Membrane Mobility

Although diffusion on the plasma membrane is typically described as two-dimensional, three-dimensional structures such as blebs^46^ and tube-like protrusions^47, 48^ can significantly compartmentalize diffusion^19^. To explore the effects of cellular topography on membrane diffusivity, we conducted concurrent 3D cellular membrane imaging via astigmatism^49^ and point-cloud diffusivity mapping. Fig. 4a and 4b illustrate the 3D and pc-SM*d*M images, revealing numerous ∼100 nm bulges on the PM (indicated by orange arrows in Fig. 4a,b), with slow diffusion rates down to ∼1 μm^2^/s. Examination of diffusivity on the ER tubule unveiled slowdowns at ER-PM contact sites (green arrows in Fig. 4a,b), aligning with previous studies^39^. These topographical heterogeneities are further depicted in Fig. 4a, showcasing the bulges on the PM and the ER-PM contact sites in the *x*-*z* plane (marked by orange and green boxes respectively). Time series of the images in the *x*-*z* plane indicate these bulges are linked to endocytosis/exocytosis (Extended Data Fig. 4a), consistent with clathrin-coated pit-induced diffusion impediments through altered membrane geometry.

**Fig. 4:**
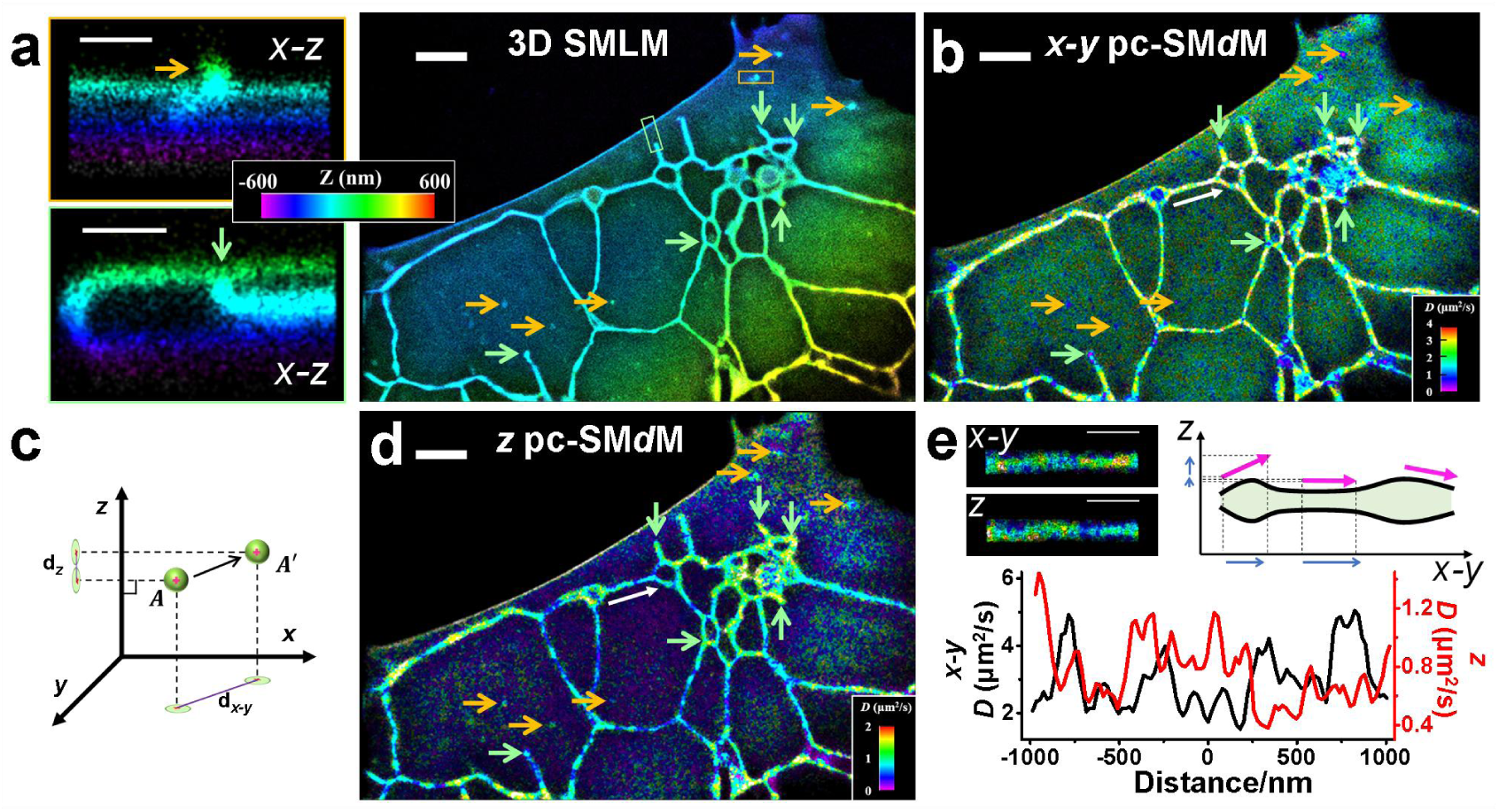
Topographic impacts on membrane diffusivity revealed by concurrent 3D-SMLM and pc-SM*d*M imaging. **a,** 3D-SMLM image of a live cell membrane using BDP-TMR-azide, highlighting nanoscale topographical features. Enlarged views of the orange boxed region show a membrane bulge, while the green boxed region indicates an ER-PM contact site. **b,** pc-SM*d*M image depicting diffusivity in the *x*-*y* plane. **c,** Projection of single-molecule displacements onto the *z*-axis, used to reconstruct the pc-SM*d*M image along the *z*-axis. **d,** pc-SM*d*M image along the *z*-axis, showing topographical features, including the PM bulge and ER-PM contact site (indicated by orange and green arrows, respectively, in panels **a**, **b**, and **d**. **e,** Correlative analysis between *x*-*y* plane and *z*-axis pc-SM*d*M images reveals a negative correlation, likely due to topographical variations of ER tubules.

The *x*-*y* plane pc-SM*d*M primarily captures diffusion rates based on displacement projections onto the *x*-*y* plane. However, diffusion in the *x*-*y* plane can be confined when the 2D membrane extends into the *z*-direction. To account for this, we generated a *z*-axis diffusivity mapping by projecting displacements onto the *z*-axis (Fig. 4c). Fig. 4d displays this *z*-axis pc-SM*d*M image, highlighting increased *z*-axis diffusion rates at the PM bulges and ER-PM contact sites (marked by orange and green arrows, respectively). Interestingly, a comparison of the diffusion rates in the *x*-*y* plane versus the *z*-axis for an ER tubule reveals a negative correlation (Fig. 4e, corresponding to white arrow in Fig. 4b,d). indicating a close link between diffusion rates and ER tubule topography. A recent study has identified two distinct forms of ER tubules with varying diameters^50^. Through three-dimensional pc-SM*d*M, we speculated that at junctions where these two forms converge, diffusion rates decrease in the *x*-*y* plane but increase in the *z*-axis (Fig. 4e). Similarly, in the *x*-*y* plane, the diffusional slowdowns observed at ER-PM contact sites are due to the extension of the ER tubule into the *z*-direction. The dynamic variation of ER diameters is further demonstrated in time-series images in the *x*-*z* plane (Extended Data Fig. 4a). Topographical alterations may also contribute to diffusivity slowdowns in the nuclear ER at the junctions between the nuclear envelope (Extended Data Fig. 4b-d) and nuclear ER, as well as in the tube-like protrusions on the PM (Extended Data Fig. 4e-g).

### Interplays Between the Diffusivity Heterogeneities of Lipids and Membrane Probes

The diffusivity of lipids, crucially influenced by their chemical structures and interactions with local membrane architectures, plays a pivotal role in regulating membrane functionalities. Our pc-SM*d*M technique, when integrated with a two-color SMLM setup, enables the concurrent and correlative analysis of diffusional heterogeneities of two distinct lipid types. As cholesterol plays a central role in regulating lateral mobility on cellular membranes, we thus explored the interplay between the diffusion heterogeneities of cholesterol and dioleoyl phosphatidyl ethanolamine (DOPE), an unsaturated lipid. Given the challenge in the uptake of fluorescently labeled cholesterol by live cells, we tagged cholesterol with a single-stranded DNA (docking strand) at its hydrophilic end^51^, facilitating its imaging utilizing DNA-PAINT^52, 53^ with a fluorescently labeled complementary sequence (imager strand). Concurrently, we monitored DOPE’s diffusion using lissamine rhodamine-labeled DOPE. The chemical structures of these lipids are detailed in Extended Data Fig.8a.

Fig. 5a and 5b illustrate concurrent pc-SM*d*M images of cholesterol and DOPE, showing the plasma membrane but not organelle membranes, confirming that the labeled lipids are predominantly anchored in the upper leaflet of the PM. Diffusivity mapping revealed nanoscale heterogeneities for both lipids, with average diffusion rates of 0.46 ± 0.06 μm^2^/s for cholesterol and 0.20 ± 0.03 μm^2^/s for DOPE (Fig. 5j).

**Fig. 5:**
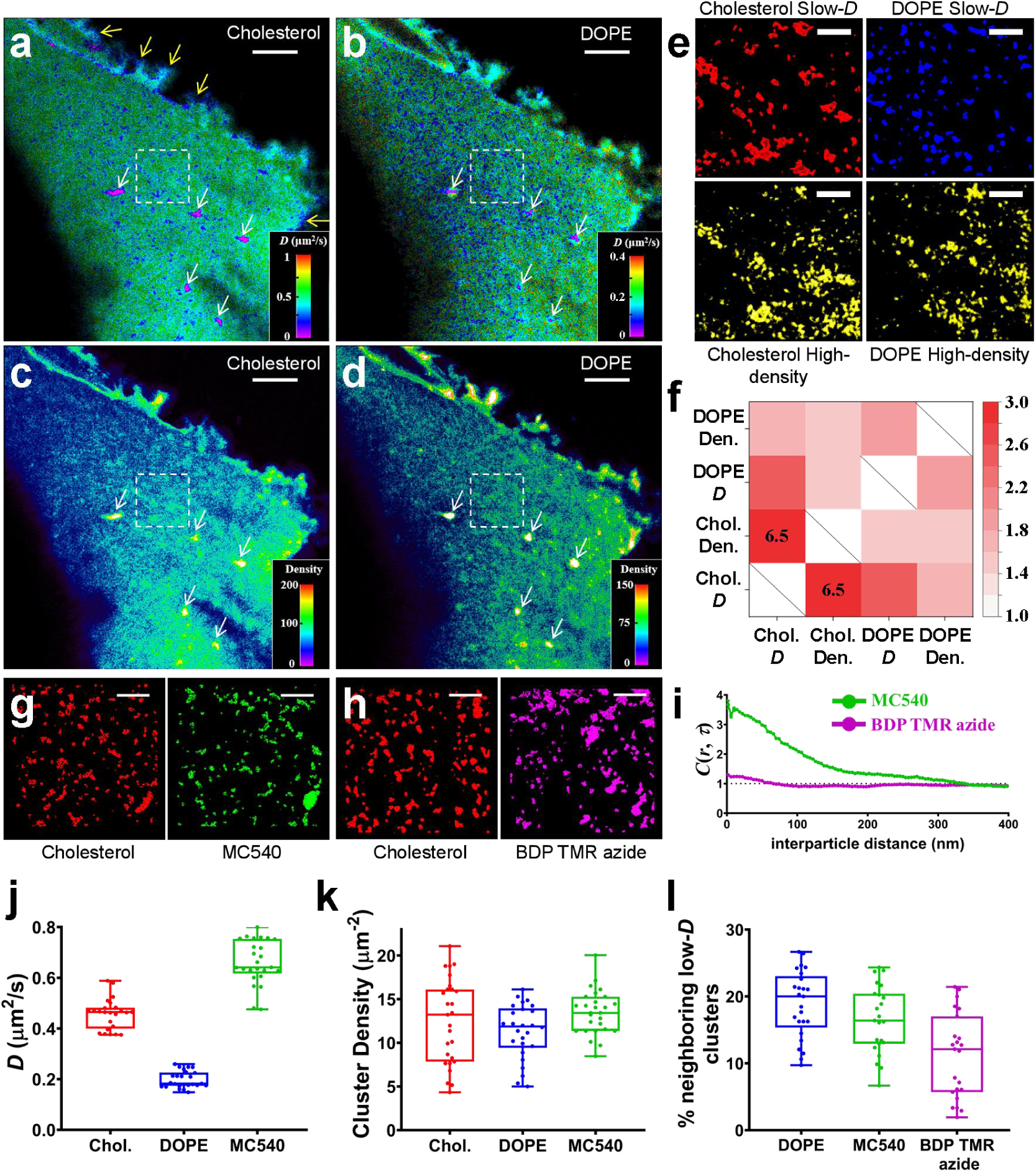
Distinct diffusion heterogeneities for multiple lipids and membrane probes revealed by concurrent two-color pc-SM*d*M imaging. **a,** pc-SM*d*M image of cholesterol on the live cell plasma membrane, showing cholesterol-rich regions that impede diffusion (white arrows). Diffusion within filopodia regions is notably slower compared to other areas (yellow arrows). **b,** Concurrent pc-SM*d*M image of DOPE. **c-d,** Density-SMLM images of cholesterol and DOPE, with color representing local single-molecule density. **e,** "Fingerprint" maps generated by extracting slow-*D* clusters from white boxes in panels **a** and **b**, and high-density clusters from white boxes in panels **c** and **d**. **f,** Heatmap showing cross-correlation analysis between pairs of images in panel **e**. **g,** Comparison of "fingerprint" maps for slow-*D* clusters of cholesterol and MC540. **h,** Comparison of "fingerprint" maps for slow-*D* clusters of cholesterol and BDP TMR azide. **i,** Cross-correlation analysis reveals stronger similarity in diffusion heterogeneities between cholesterol and MC540 than between cholesterol and BDP TMR azide. **j,** Average diffusion rates for cholesterol, DOPE, and MC540 on the live cell plasma membrane, derived from analyses across 25-30 regions in 10 cells. **k,** Density of slow-*D* clusters for cholesterol, DOPE, and MC540 on the plasma membrane, derived from analyses across 25-30 regions in 10 cells. **l,** Likelihood of finding a neighboring slow-*D* cholesterol cluster for DOPE, MC540, and BDP TMR azide clusters, derived from analyses across 25-30 regions in 8-10 cells. Scale bar: 2 μm in **a**, **b**, **c**, and **d**, 500 nm in **e**, **g**, and **h**.

Cholesterol’s diffusion rate is consistent with measurements from a smaller molecular weight fluorescently-labeled cholesterol (Extended Data Fig.5) and previously reported values^54^. Corresponding Density-SMLM images (Fig. 5c,d) indicate local single-molecular density, with large cholesterol-rich clusters impeding diffusion for both cholesterol and DOPE (white arrows, Fig. 5a-d).

In zoomed-in regions without large cholesterol-rich clusters (white boxes, Fig. 5a-d), we extracted slow-*D* and high-density clusters to construct characteristic maps for cholesterol and DOPE (Fig. 5e), generating "fingerprint"-like patterns to investigate their spatial correlations. Cross-correlation analysis of these maps produced a heatmap (Fig. 5f), revealing strong correlations between cholesterol slow-*D* clusters and cholesterol high-density clusters, as well as between cholesterol slow-*D* clusters and DOPE slow-*D* clusters. These findings suggest that local cholesterol concentration predominantly dictates the diffusivity of both cholesterol and unsaturated lipids. Additionally, a moderate correlation was observed between cholesterol slow-*D* clusters and DOPE high-density clusters, indicating a slight enrichment of unsaturated lipids within cholesterol slow-*D* clusters.

In filopodia regions (yellow arrows, Fig. 5a), cholesterol diffusion was notably slower than DOPE (Extended Data Fig.6b-d). Cholesterol enrichment in filopodia is known to facilitate the formation of narrow, finger-like protrusions by enhancing membrane flexibility^55, 56^. Moreover, recent studies suggest the presence of mobile cholesterol, distinct from lipid raft-associated cholesterol, in filopodia^57^. Our diffusivity mapping may support the hypothesis that cholesterol plays a critical role in signaling pathways related to filopodia formation and extension, potentially through interactions with actin-associated membrane proteins.

We further explored the correlation between cholesterol and other membrane probes by constructing diffusivity fingerprint maps. Fig. 5g and 5h show zoomed-in images of cholesterol with merocyanine 540 (MC540) and BDP TMR azide, respectively (full images in Extended Data Fig.7a,b). Cross-correlation analysis revealed that MC540’s diffusional behavior closely resembled that of cholesterol, while BDP TMR azide exhibited different behavior (Fig. 5i and Extended Data Fig.7c).

These differences arise from the orientation of the probes within the lipid bilayer, governed by their hydrophobic interactions with the bilayer. MC540, a amphiphilic membrane probe, is horizontally anchored within the bilayer, with its charged sulfonic group exposed to anions and water^58^ (Extended Data Fig.8b), a configuration akin to that of amphiphilic lipids and cholesterol^59^. Conversely, BDP TMR azide, a neutral, cell-permeable molecule, diffuses freely between lipid layers and targets intracellular organelles. Its flexible orientation^54, 60^ contributes to its differing diffusion behavior (Extended Data Fig.8b). The density of the slow-*D* clusters of these membrane probes were presented in Fig. 5k. Notably, cholesterol clusters exhibited slightly higher density and smaller diameters compared to DOPE clusters (Extended Data Fig. 7d-f). Additionally, we quantified the likelihood of encountering a neighboring slow-*D* cholesterol cluster for DOPE, MC540, and BDP TMR azide, demonstrating stronger correlations with cholesterol for DOPE and MC540 than for BDP TMR azide (Fig. 5l). These findings highlight the potential of diffusivity fingerprint maps as a tool for identifying diffusers with shared molecular interactions within membranes.

## Discussion

By developing point-cloud single-molecule diffusivity mapping (pc-SM*d*M), we achieved a spatial resolution of ∼50 nm for mapping membrane mobility in live cells. This enhanced resolution allowed us to directly visualize nanoscale slow-*D* clusters (clusters with diffusion slowdowns) on the plasma membrane, resembling the ’pickets’ that impede membrane mobility. While this study primarily focused on slow diffusion (∼1 μm^2^/s) on cellular membranes, our algorithm is versatile and can be readily adapted to analyze fast diffusion in the live cell cytosol and nucleus (Extended Data Fig.9). This capability enables detailed insights into diffusion slowdowns caused by cytoskeletal structures, such as actin stress fibers and chromatin. Furthermore, the same Gaussian denoising filter-based approach can be seamlessly extended to other multi-dimensional super-resolution imaging techniques, particularly those reliant on statistical analysis to extract physicochemical properties.

The lateral mobility of the plasma membrane is influenced by several interconnected factors, including lipid composition, protein assemblies, and membrane topography. Previous studies have often examined these factors in isolation, relying on ensemble measurements to infer their effects. Using our point-cloud super-resolution diffusivity mapping coupled with correlative super-resolution imaging techniques, we revealed that diffusion slowdowns on the plasma membrane are highly heterogeneous and result from complex molecular interactions. Cholesterol content emerged as a dominant regulator, while cytoskeleton-associated protein assemblies contributed to diffusion confinement in only a few localized spots. Additionally, membrane topographical features such as blebs and protrusions were found to compartmentalize diffusion in the *x*-*y* plane, acting as ’bumps’ that hinder lateral mobility. This approach uniquely enables the direct spatial correlation of membrane diffusivity heterogeneities with structural heterogeneities on cellular membranes.

Beyond nonspecific interactions affecting all diffusers, specific lipid-protein interactions, driven by the chemical structures of lipids, introduce an additional layer of complexity to diffusional regulation. These interactions, such as the clustering of signaling proteins, generate ’diffusional traps’ that concentrate signaling molecules, thereby enhancing signal transduction within regions of local lipid composition heterogeneities. Using our two-color diffusivity mapping technique, we successfully distinguished the diffusional heterogeneities among different lipids. Notably, cholesterol exhibited significantly slower diffusion rates in filopodia compared to DOPE, highlighting its pivotal role in filopodia formation and extension. The high-density and slow-*D* clusters of cholesterol observed on the plasma membrane are likely stable lipid aggregations enriched with cholesterol and unsaturated lipids, persisting over minutes. This stands in contrast to the transient nature of lipid rafts, which are thought to exist on the timescale of milliseconds to seconds and are predominantly composed of cholesterol and saturated lipids. Moving forward, elucidating the unique diffusional heterogeneities of various lipids and membrane proteins, along with identifying their interacting partners using diffusivity fingerprint maps, remains an exciting and critical direction for advancing the pc-SM*d*M technique.

## Conclusions

In conclusion, we have developed point-cloud single-molecule diffusivity mapping (pc-SM*d*M), achieving a significant advancement in the spatial resolution of diffusivity mapping on live cell membranes, reaching ∼50 nm. This innovation allows for the direct visualization of nanoscale slow-*D* clusters while maintaining the fidelity of point-cloud data. By providing high-resolution insights into molecular interactions and leveraging correlative super-resolution microscopy (SMLM), we revealed how lipid composition, membrane-associated proteins, and cellular topography collectively influence lateral mobility on plasma membranes. Through two-color diffusivity mapping, we uncovered distinct fingerprint maps of diffusion heterogeneities among various lipids and membrane probes, determined by their interactions with lipid bilayers. Notably, cholesterol emerged as a unique factor, significantly slowing diffusion in filopodia and highlighting its role in modulating membrane flexibility and facilitating signaling pathways. Overall, pc-SM*d*M offers a powerful tool for high-resolution studies of molecular diffusion in live cells, paving the way for deeper exploration of nanoscale heterogeneities in biological membranes and the molecular mechanisms underlying membrane dynamics.

## Methods

### Cell culture and sample preparation

The COS-7 cell line (Procell, CL-0069) was purchased from Wuhan Pricella Biotechnology Co., Ltd. The cells were cultured in a T25 cell culture flask (Corning) using DMEM media supplemented with 10% fetal bovine serum and 1% penicillin-streptomycin, maintained in a 5% CO_2_ atmosphere at 37°C. Two days before imaging, the cells were plated onto 18-mm diameter glass coverslips (CITOGLAS, 18 mm diameter, #1.0H) that had been pretreated with hot piranha solution (H_2_SO_4_: 30% H_2_O_2_ at 3:1). Prior to imaging, the coverslip was transferred to a holder (Bioscience Tools, CSC-18) compatible with the microscope stage.

### Plasmid constructs and transfection

mEos3.2-C1 was a gift from Michael Davidson and Tao Xu (Addgene plasmid no. 54550) and was used without modification. mEos3.2-NLS was constructed by inserting the desired DNA sequence (GENEWIZ Co. Ltd.) between the SacⅠ and BamHI restriction enzyme recognition sites within the short sequence at the C terminus of mEos3.2-C1. Verification of plasmid constructs was confirmed through Sanger sequencing. Cells were allowed to grow up to ∼60% confluency before being transfected with the Lipofectamine 3000 (ThermoFisher) according to the recommended protocol. The following sequences of plasmid were used in this study:

mEos3.2-C1: mEos3.2-SGLRSRAQASNSAVDGTAGPGSTGSR

mEos3.2-NLS: mEos3.2-SGLRSRADPKKKRKVDPKKKRKVDPKKKRKVGSTG SR

### Fluorescent Probes

BDP-TMR-alkyne and BDP-TMR-azide were purchased from Aladdin (B171327 and B171329). Merocyanin 540 was purchased from bidepharm (BD01090609). 18:1 Liss Rhod PE (DOPE) was purchased from Avanti Polar Lipids (810150P).

### Fluorescent Labelling

For pc-SM*d*M imaging of live cells, the samples were prepared in an imaging buffer consisting of Leibovitz’s L-15 medium (Procell, PM151013) supplemented with 20 mM HEPES (Beyotime, C0215). BDP-TMR-alkyne, BDP-TMR-azide, Merocyanin 540, or 18:1 Liss Rhod PE were diluted in the imaging medium to a final concentration of 1-3.3 nM. The dye-containing imaging medium was directly added to the samples, followed by the image collections.

For SMLM imaging of Clathrin, Ankyrin, and Adducin, the samples were chemically fixed using 4% paraformaldehyde for 10 min and were washed twice with phosphate-buffered saline (PBS) buffer. The cells were then blocked in a blocking buffer (3% bovine serum albumin + 0.05% Triton-X in PBS) and incubated for 1 hour at room temperature with primary antibodies. After three washes in washing buffer (0.2% bovine serum albumin in PBS), the cells were incubated for 45 minutes at room temperature with secondary antibodies. The samples were washed with washing buffer for 5 minutes, three times before imaging in a photoswitching buffer (PBS containing 5% glucose, 200 mM cysteamine, 0.8 mg/mL glucose oxidase, and 40 µg/mL catalase). The primary antibodies used were rabbit anti-Clathrin light chain (Abcam, ab271185), mouse anti-ANK2 (Santa Cruz Biotech, sc-12718), and mouse anti-Adducin α (Santa Cruz Biotech, sc-33633). The secondary antibodies used were goat anti-rabbit and goat anti-mouse, conjugated with Alexa Fluor 647 (Abcam, ab150087 and ab150119) at a dilution of 1:400.

### Cholesterol depletion and addition treatment

Live cells were chemically fixed using a solution of 3% paraformaldehyde and 0.1% glutaraldehyde in PBS followed by two washes with 0.1% sodium borohydride in PBS. Fixed cells were treated with 5 mM solution of methyl-β-cyclodextrin (MβCD; Beyotime, ST1515) and 5 mM solutions of water-soluble cholesterol (cholesterol-MβCD; Sigma, C4951) in PBS for 15∼30 min, respectively. Cells were then gently washed twice with DPBS.

### CTB treatment

Cells were incubated with 1 μg/mL Alexa Fluor 647-conjugated CTB (Invitrogen, C34778) in the culture medium for 5 min at room temperature, and then washed twice with the imaging buffer before imaging.

### DNA-PAINT experiments for cholesterol imaging

Live cells were treated with a 5 mM solution of MβCD in the culture medium for 20 minutes in 5% CO_2_ at 37°C, followed by two washes with imaging buffer. The sample was then incubated with a docking strand (500 nM) for 10 minutes at room temperature, and then washed with imaging buffer. A final concentration of 2 nM imager strand was added to the imaging medium for imaging. The following DNA sequences are used in this study:

Docking strand: cholesterol - TEG - conjugated:

cholesterol-TEG-TCTCTCTCTCTCTCTCTCTCTCTCTCTCTCTCTCTCTCTCTCTCTCTCTCTCTCTCTCTCTCTCTCTCTCTCTCT, containing 37.5×TC.

imager strand: AF647 - conjugated: 4×(GA)-AF647

GAGAGAGA-AF647

### Optical setup

SMLM, 3D-SMLM, pc-SM*d*M and two color pc-SM*d*M were performed on a home-built setup based on an Olympus IX83 inverted fluorescence microscope. Briefly, 405 nm (CNIlaser, MDL-III-405, 500 mW), 488 nm (OBIS 488 LX, Coherent, 150 mW), 532 nm (CNIlaser, OEM-U-532, 500 mW), 560 nm (MPB Communications, 500 mW) and 642 nm (MPB Communications, 500 mW) lasers were focused at the back focal plane of an oil-immersion objective lens (Olympus UPLXAPO 100X, NA 1.45) to enter the coverslip–sample interface slightly below the critical angle, thus illuminating a few micrometers into the cell.

### pc-SM*d*M with BDP-TMR-azide

For pc-SM*d*M with BDP-TMR-azide, the sCMOS camera (Teledyne Photometrics, Prime 95B) was used in effective global exposure mode and synchronized with the laser, recording continuously at ∼140 fps, with the 532 nm lasers outputting tandem pulses. The typical center-to-center separation between the paired pulses was Δt ≈ 6 ms (Fig. 1a). The estimated peak and average power densities at the sample were ∼1.4 and ∼0.4 kW/cm^2^, respectively. Wide-field fluorescence emission was filtered by a long-pass filter (Chroma, ET542lp) and a band-pass filter (400-630 nm). 60,000-80,000 frames of single-molecule images were recorded, accumulating ∼10⁶ molecules across the view.

### Concurrent pc-SM*d*M and 3D-SMLM

For 3D localization, a cylindrical lens was used to induce elongations of single-molecule images in vertical and horizontal directions for molecules below and above the focal plane, respectively. Frame-synchronized stroboscopic excitation was achieved through direct power modulation of the 532 nm laser using the ’All Rows’ trigger mode in a global acquisition state (Teledyne Photometrics, Prime 95B). The illuminated area was ∼90 μm in diameter, yielding ∼1.2 kW/cm^2^ peak power density and ∼0.3 kW/cm^2^ average power density. Single-molecule images were recorded in wide field with the sCMOS camera at a frequency of ∼145 Hz and an effective exposure time of 2 ms. Paired pulses were applied across tandem camera frames (Fig. 1a). 65,000 frames of single-molecule images were recorded, accumulating ∼10⁶ molecules across the view.

### Concurrent two-color pc-SM*d*M

Concurrent two-color pc-SM*d*M of 18:1 Liss Rhod PE and DNA-PAINT were achieved on a custom-built system. The power of the 560 nm (for excitation of 18:1 Liss Rhod PE) and 642 nm (for excitation of the imager strand) lasers was controlled via an AOTF (Gooch & Housego, 80-152 MHz Vert.Pol.4X). The excitation beam was directed towards the objective by a four-color notch dichroic mirror (Chroma, ZT405/488/561/640rpcv2). In a 4f-system, the fluorescence emission was split by a dichroic mirror (Chroma, ZT640rdc-UF1). Fluorescence in green channel (excited by 561 nm laser) and in red channel (excited by 642 nm laser) were filtered by Chroma ET595/44m filter and Chroma ET705/100m filter respectively. Images were collected on two sCMOS cameras (Tucsen, Dhyana 95V2). The imaging setup resulted in an effective pixel size of 110 nm. Dual-color-SMLM data was recorded with our custom-built setup operating in dual-color alternate illumination mode using a multifunction I/O board (National Instruments, PCIe-6323), to reach a frame-to-frame separation time of 14.76 ms, with an acquisition frequency of 67 Hz. Experiments were performed at laser excitation powers of 20 mW for 560 nm and 25 mW for 642 nm, translating to irradiances of 0.3 kW/cm^2^ and 0.4 kW/cm^2^ peak power density, respectively. 80,000 frames of single-molecule images were recorded, accumulating ∼10⁶ molecules across the view.

### SMLM with Clathrin, Ankyrin, and Adducin

The samples were excited with stroboscopic pulses of 2 ms duration at ∼0.7 kW/cm^2^ peak power density and ∼0.2 kW/cm^2^ average power density. Wide-field emission was filtered by a long-pass filter (Chroma, ET655lp) and a band-pass filter (Chroma, ET705/100m), and recorded with the sCMOS camera at 140 fps. The sCMOS camera was synchronized with the lasers to reach an effective global exposure. 70,000 frames were recorded, and the accumulated single-molecule images were reconstructed to obtain the SMLM images.

### Data analysis for pc-SM*d*M

Single-molecule images were fitted with a 2D Gaussian function, and the centroid positions of all single molecules were accumulated and reconstructed to generate the SMLM images. SM*d*M analysis was processed as described previously. Briefly, the positions of the molecules identified in the latter frame were used to search for matching molecules in the former frame within a cutoff displacement threshold (*R*). Typical values of *R* for fast- and slow-moving molecules were generally set to be 5 pixels and 3 pixels, respectively (with a pixel size of ∼110 nm). The 2D displacements (denoted as *d*) in the x-y direction were calculated for the matched molecules. This process was repeated for all paired frames, and each point in the reconstructed super-resolution image was assigned a *d* value (the displacement).

Next, as illustrated in the main text and Fig. 1b, each single molecule in the super-resolution image served as the center of a circle in turn (the targeted molecule). All single molecules surrounding this targeted molecule within a radius of ∼50 nm were considered to generate the displacement histogram. The weighted values (*f*(*l*), the count in the displacement histogram) for all single molecules within this radius were calculated based on their distance from the central molecule (denoted as *l*), according to the following equation:

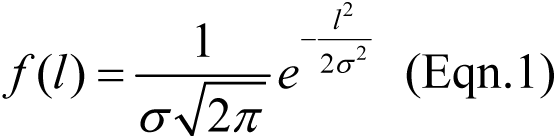

Where *l* is the distance of each single molecule from the central single molecule. The standard deviation *σ* = 42 nm is used (when search radius is 50 nm). Finally, the displacement histograms were fitted with an equation based on a modified isotropic 2D random-walk model to determine the local diffusion rate (*D*):

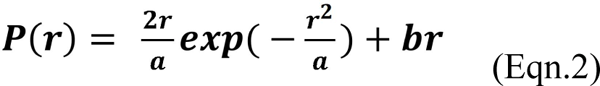

where *r* is the single-molecule displacement in the time interval Δ*t*, *a* = 4*D*Δ*t*, and *b* is a background term to account for molecules that randomly enter the view, as rationalized and validated previously with experiments carried out at different single-molecule densities. This *D* value was then assigned to the central single molecule, and the color in the diffusivity-resolved pc-SM*d*M images was determined by this *D* value.

For the analysis of *D* in the axial (*z*) direction, as shown in Fig.4, we only consider the projections of the displacement along the *z*-axis to construct the displacement histogram. For the analysis of one-dimensional motion, such as displacement along an ER tubule, we first performed principal component analysis (PCA) on all displacements to determine the preferred angle *θ*. The single-molecule displacements were then projected onto either the *z* direction (for *z* direction analysis) or the preferred angle *θ* (for one-dimensional motion along the ER tubule). The displacement histograms were fitted with a modified 1D random-walk model:

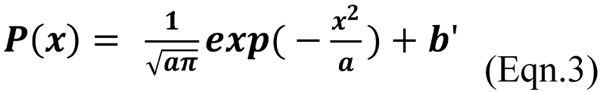

where *x* is the one-dimensional single-molecule displacement in the time interval Δ*t*, *a* = 4*D*Δ*t*. The one-dimensional displacement histograms were fitted to give the one-dimensional *D* values.

### Quantification of proteins domains and slow-*D* clusters on plasma membrane

To define the slow-*D* clusters, a Gaussian fit was applied to the *D* values of all single molecules on the plasma membrane to determine the mean value and full width at half maximum (FWHM). Slow diffusion (*D*_slow_) was defined as diffusion rates lower than the FWHM range (Extended Data Fig. 1a). Clustering analysis was performed by density-based spatial clustering of applications with noise (DBSCAN). The algorithm uses two user-selectable parameters, the search radius and the minimal point of single-molecule localizations within that radius, to identify clusters with dense single molecules. For localizing the slow diffusion rate clusters on the plasma membrane, we used 25 nm as the search radius, and less than 15 points (within the 25 nm search radius) were excluded during the analysis. The percentage of neighboring clusters was calculated by applying a 60 nm distance threshold between the two sets of cluster data.

## Acknowledgments

L.X. acknowledges financial supports from National Key R&D Program of China (2022YFA1305400), National Natural Science Foundation of China (22104113, 22274122), Fundamental Research Funds for the Central Universities interdisciplinary (2042023kf1012), and Innovative Talents Foundation from Renmin Hospital of Wuhan University (JCRCFZ-2022-010). S. J. acknowledges financial support from National Natural Science Foundation of China (22302092).

## Author contributions

C. H. and L. X. conceptualized and designed the study. C. H. conducted all cell imaging experiments. Z. Z. and L. X. developed and constructed the imaging system. H. G. contributed to the pc-SM*d*M programming and calibration for two-color pc-SM*d*M imaging. J. D. and Q. W. assisted in designing the DNA-PAINT experiments. Y. W. and L. L. carried out the calibration and data analysis for DBSCAN. Y. W. and L. X. (Lin Xu) performed the calibration for 3D-STORM. S. J. supervised the DNA-PAINT imaging work. X. Z. and W. H. provided input during discussions and validation of the pc-SM*d*M program. K. C., R. Y., W. L., and K. X. reviewed and edited the manuscript. K. X. and L. X. supervised the entire study. C. H. and L. X. wrote the manuscript with contributions from all authors.

## Data availability

The data that support the findings of this study are available from the corresponding author upon reasonable request.

