## Supplementary Figures for "Super-Resolution Diffusivity Mapping Reveals Spatial Correlations Between Lateral Mobility and Structural Heterogeneities on Cellular Membranes"

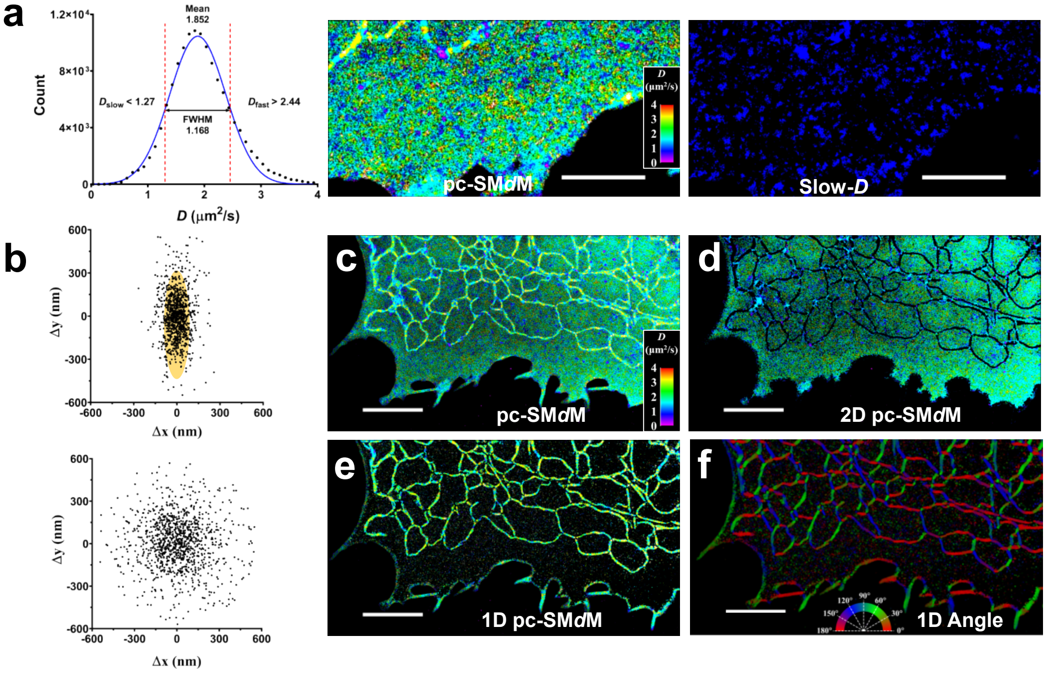


**Extended Data Fig. 1: Data processing and analysis of point-cloud single molecule diffusivity mapping (pc-SM*d*M).**

**a,** Identification and extraction of slow-*D* clusters based on the distribution of single-molecule diffusion rates on the plasma membrane. A Gaussian fit was applied to the distribution of diffusion rates (*D* values) for all single molecules on the plasma membrane, yielding the mean value and the full width at half maximum (FWHM). Molecules with diffusion rates below the FWHM range were categorized as slow-diffusion ($D_{slow}$), while those with rates above the FWHM range were categorized as fast-diffusion ($D_{fast}$). Using this classification, images of slow-*D* clusters were generated from the point-cloud single-molecule diffusivity mapping data, enabling precise spatial visualization of slow-diffusion regions. **b,** 2D plot of single-molecule displacements along an ER tubule (top) and on the plasma membrane (PM) (bottom). Diffusion is considered one-dimensional if the anisotropy value exceeds 0.6 (top). **c-f,** pc-SM*d*M image displaying both one-dimensional and two-dimensional diffusion, only two-dimensional diffusion, only one-dimensional diffusion, and angle image for one-dimensional diffusion extracted from PCA with anisotropy values larger than 0.6. Scale bar: 2 μm.

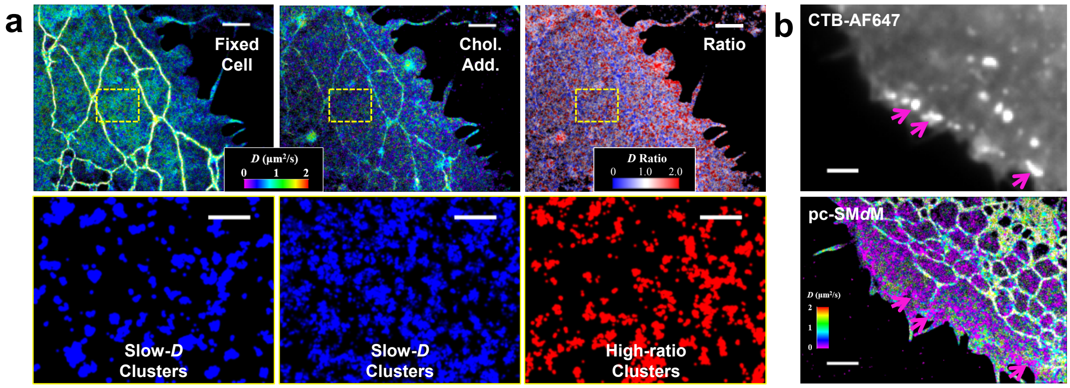


**Extended Data Fig. 2: Effects of cholesterol addition and Cholera Toxin Subunit B (CTB) treatments on membrane diffusivity.**

**a,** pc-SM*d*M images of a fixed cell before and after cholesterol addition. Cholesterol addition induced more slow-*D* clusters on the plasma membrane. **b,** CTB treatment in a live cell induced the formation of nanodomains on the plasma membrane, leading to local diffusion slowdowns (indicated by magenta arrows). Top: epifluorescence image of AF647-labeled CTB; Bottom: pc-SM*d*M image. Scale bar: 2 μm for **a** and **b**; 500 nm for the zoom-in of panel **a**.

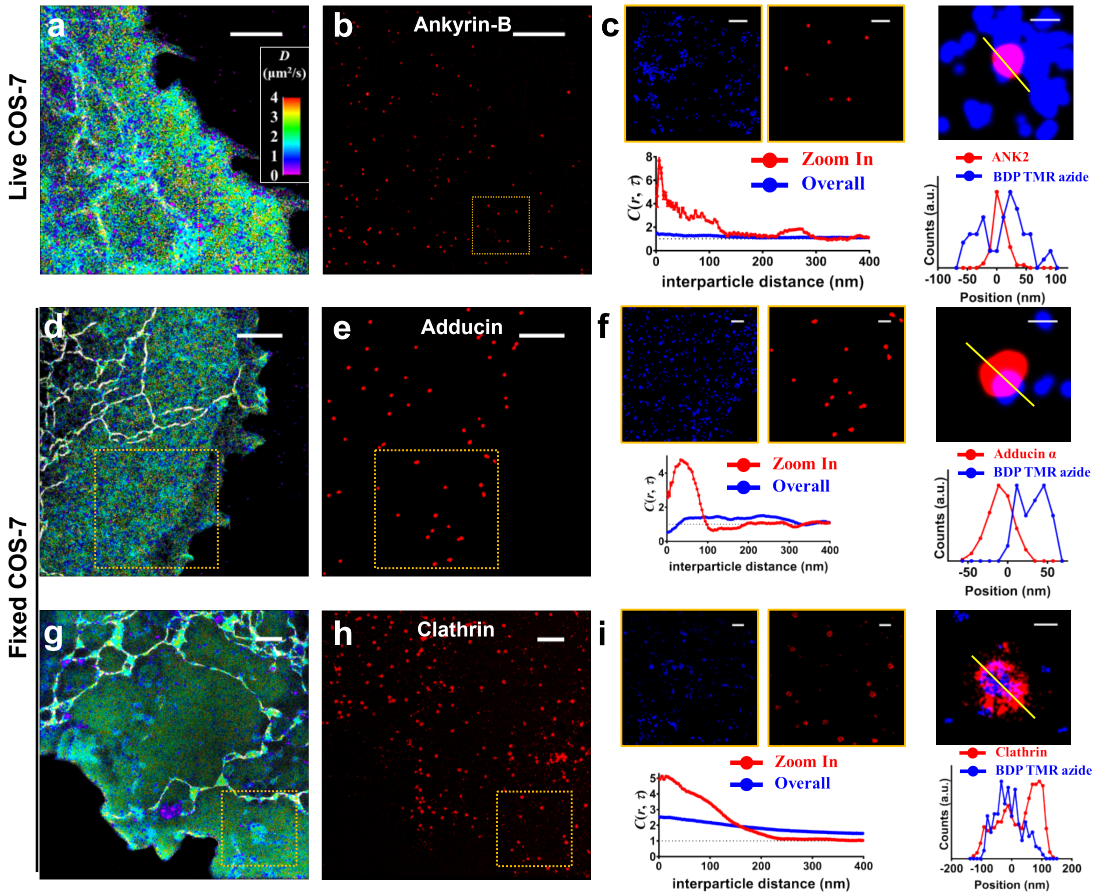


**Extended Data Fig. 3: Membrane-associated proteins partially impede local diffusion on the plasma membrane of cells.**

**a-b,** Correlative pc-SM*d*M image of a live cell and SMLM image of ankyrin-B. Scale bar: 2 µm. **c,** Zoom-in areas (orange boxes in **a** and **b**) and cross-correlation curves for the entire image (blue) and the zoom-in area (red). Cross-sectional profile of ankyrin-B merged with a slow-*D* cluster reveals a similar pattern to adducin. Scale bar: 500 nm for zoom-ins and 100 nm for cross-section. **d-e,** Correlative pc-SM*d*M and SMLM images of adducin-α in a fixed cell. Scale bar: 2 µm. **f,** Zoom-in areas (orange boxes in **d** and **e**) with cross-correlation curves. Scale bar: 500 nm for zoom-ins and 100 nm for cross-section. **g-h,** Correlative pc-SM*d*M and SMLM images of clathrin in a fixed cell. Scale bar: 2 µm. **i,** Zoom-in areas (orange boxes in **g** and **h**) with cross-correlation curves. Scale bar: 500 nm for zoom-ins and 100 nm for cross-section.

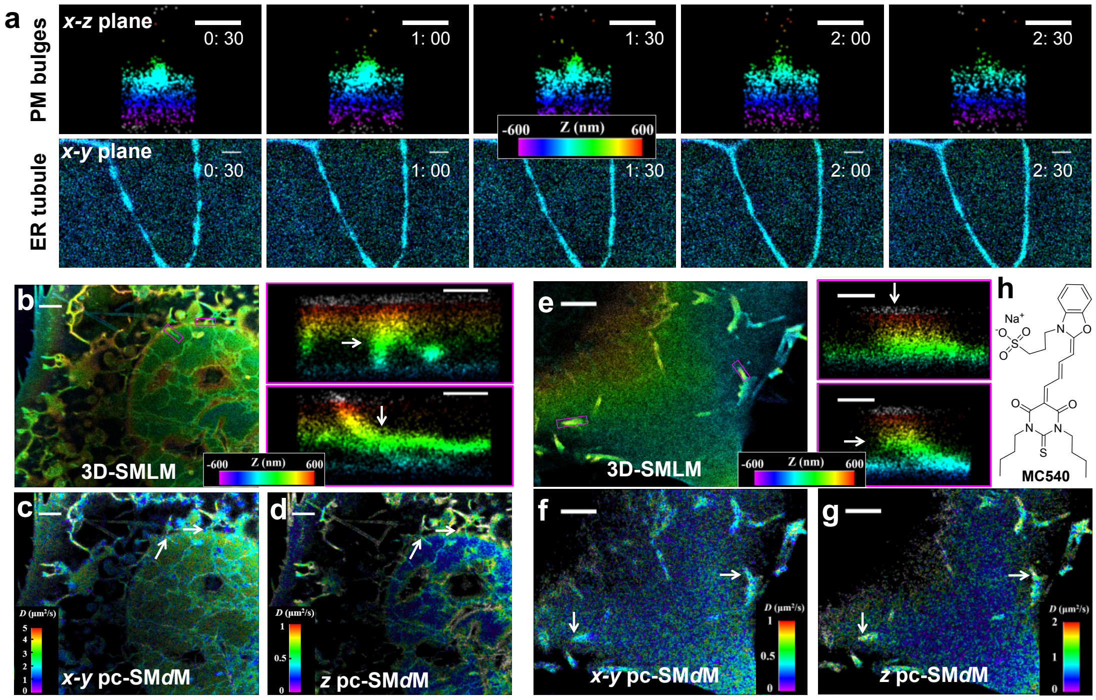


**Extended Data Fig. 4: Impacts of three-dimensional topography on membrane diffusivity.**

**a,** 3D-SMLM monitored endocytosis/exocytosis on live cell PM and dynamic contraction/expansion of ER tubules. Top: time-series images of PM bulges in the *x-z* plane. Bottom: time-series images of ER tubule in the *x-y* plane. Scale bar: 500 nm. **b,** 3D-SMLM image of a live cell's nuclear and ER membrane, with zoom-in views highlighting the junctions between the nuclear membrane and nuclear ER (magenta boxes). Scale bar: 2 µm; 500 nm for zoom-in. **c-d,** pc-SM*d*M image of the *x-y* plane and along the *z*-axis. Diffusion rates are slower on the *x-y* plane but faster along the *z*-axis (white arrows), reflecting changes in the dimensionality of diffusion at these structural junctions. Scale bar: 2 µm. **e,** 3D-SMLM image of the plasma membrane with Merocyanine 540 (MC540). Zoom-in images show membrane protrusions (magenta box). Scale bar: 2 µm; 500 nm for zoom-in. **f-g,** pc-SM*d*M image in the *x-y* plane and along the *z*-axis. Protrusions slow diffusion in the *x-y* plane but increase diffusion in the *z*-axis (white arrows), revealing the impacts of dimensionality on measured diffusion rates. Scale bar: 2 µm. **h,** Chemical structure of MC540.

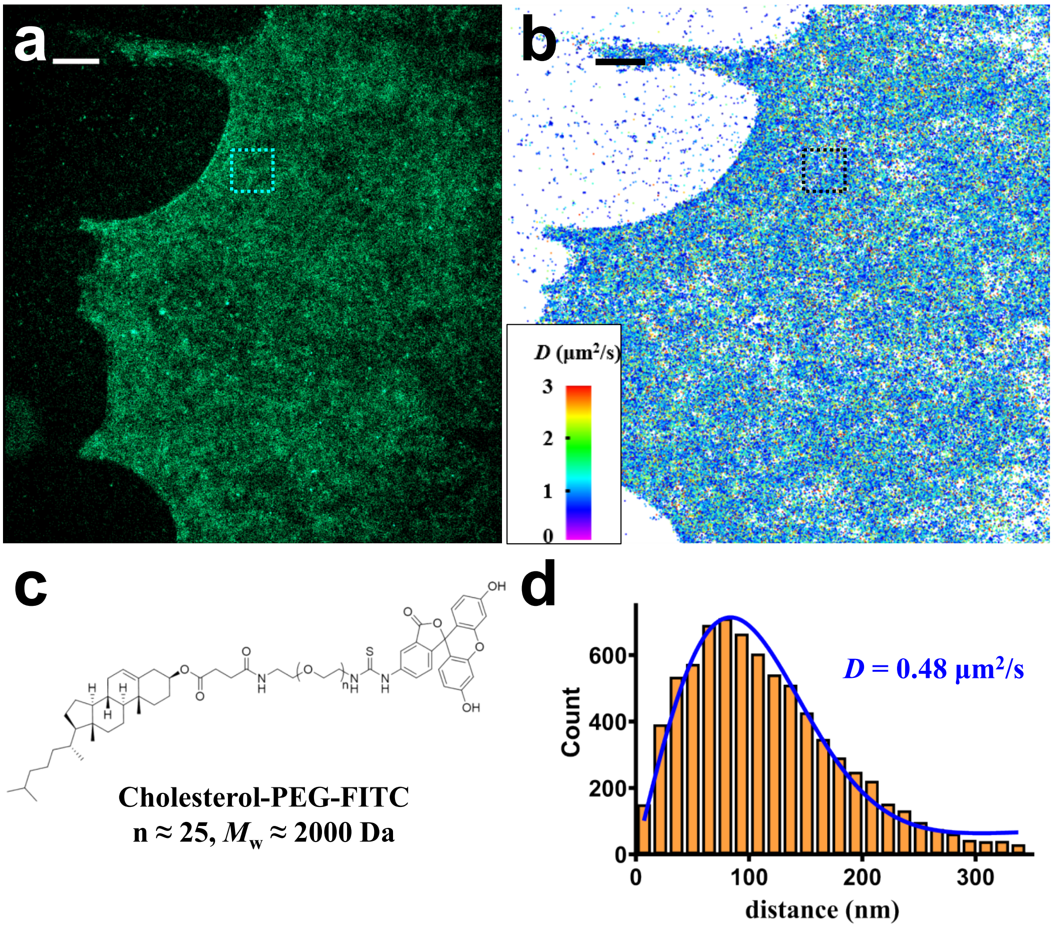


**Extended Data Fig. 5: pc-SM*d*M of cholesterol-PEG-FITC on the plasma membrane of a live COS-7 cell.**

**a,** SMLM image of the plasma membrane. **b,** Corresponding pc-SM*d*M image. **c,** Chemical structure of cholesterol-PEG-FITC. **d,** Displacement distribution (*d*) for the cyan box in panel a and black box in panel b. Fitting yielded an average diffusion rate of 0.48 μm²/s, consistent with results from the DNA-modified cholesterol probe. However, due to cholesterol-PEG-FITC's limited ability to integrate into the plasma membrane and compensate for photobleaching loss, the low throughput of single-molecule displacements resulted in the poor spatial resolution seen in panel **b**. Scale bar: 2 μm.

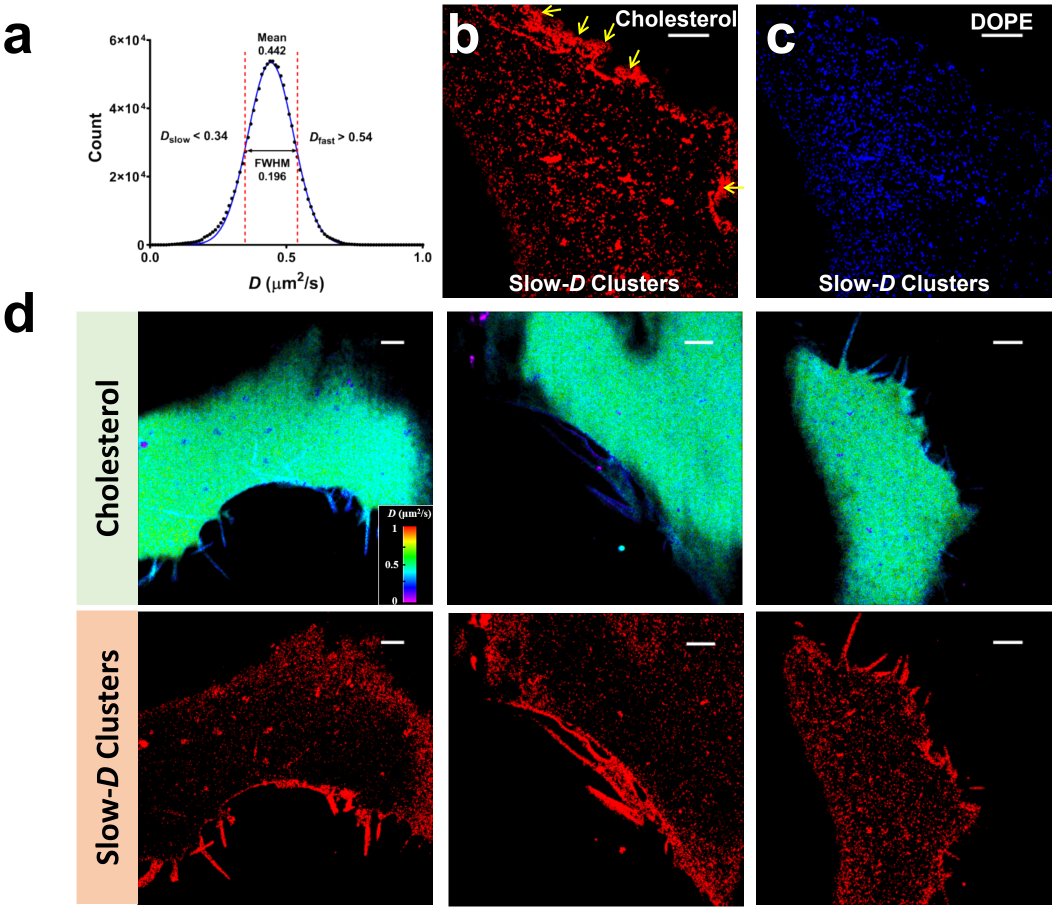


**Extended Data Fig. 6: Distinct diffusivity heterogeneities for cholesterol and DOPE on live cellular plasma membrane.**

**a,** Similar to Extended Data Fig. 1, for cholesterol and DOPE, a Gaussian fit was applied to the *D* values of all single molecules on the plasma membrane to determine the mean value and full width at half maximum (FWHM). Slow diffusion ($D_{\text{slow}}$) was defined as diffusion rates lower than the FWHM range, while fast diffusion ($D_{\text{fast}}$) was defined as those higher than the FWHM range. **b-c,** Point-cloud image of slow-*D* cholesterol clusters and DOPE clusters, extracted from Fig.5a and 5b respectively. Cholesterol showed significantly slower diffusion in the filopodia compared to other regions (yellow arrows in **b** and Fig.5a). In contrast, no similar slowdown was observed for DOPE in the filopodia. **d.** Additional slow-*D* cluster images of cholesterol on the plasma membrane of live cells, demonstrating the slower diffusion rates of cholesterol in the filopodia regions. Scale bar: 2 μm.

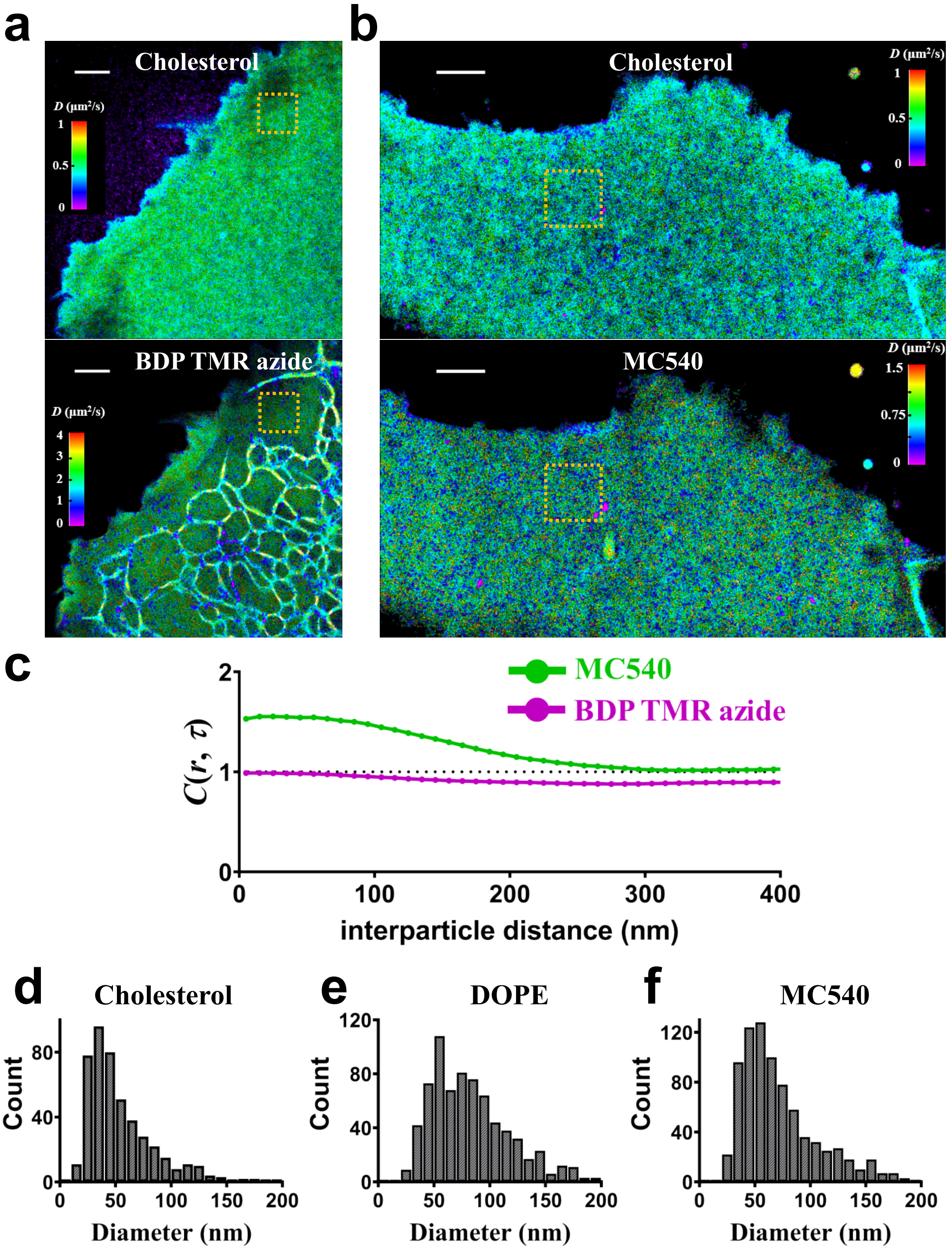


**Extended Data Fig. 7: Two-color pc-SM*d*M images and correlation analysis for cholesterol and BDP TMR azide, cholesterol and MC540.**

**a,** Concurrent two-color pc-SM*d*M imaging of cholesterol (top) and BDP TMR azide (bottom). Zoom-in images for the orange boxes are displayed in Fig.5e. **b,** Concurrent two-color pc-SM*d*M imaging of cholesterol (top) and MC540 (bottom). Zoom-in images for the orange boxes are displayed in Fig.5d. **c,** Cross-correlation analysis of slow diffusion clusters for the entire images, showing greater similarity between cholesterol and MC540 compared to BDP TMR azide. **d-f,** Size distribution of slow-*D* clusters for cholesterol, DOPE, and MC540, with DOPE showing relatively larger cluster sizes. Scale bar: 2 μm.

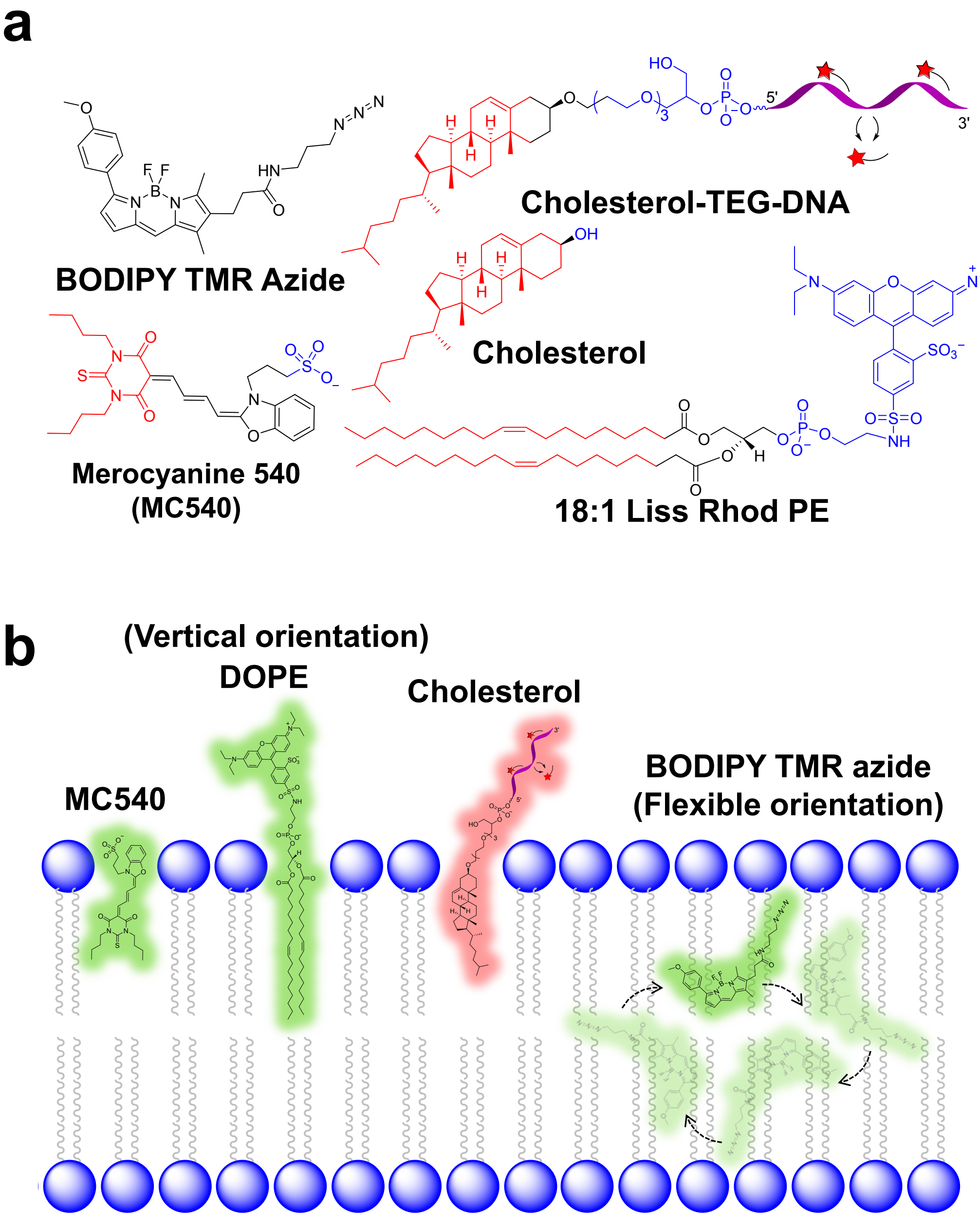


**Extended Data Fig. 8: The orientation of lipid and membrane probes on the cell membrane influences their diffusion rates.**

**a,** Chemical structures of the lipid and membrane probes used in this study. The blue region represents the charged or hydrophilic portion of the molecule, while the red section is embedded within the phospholipid membrane. **b,** BDP TMR azide, being neutral, lipophilic, and membrane-permeable, adopts a more flexible orientation within the phospholipid bilayer. In contrast, cholesterol, MC540, and DOPE are vertically oriented in the membrane, owing to their hydrophilic or negatively charged groups.

This orientation limits their ability to permeate the membrane and stain intracellular organelles. The colors of the molecules correspond to their respective excitation wavelengths.

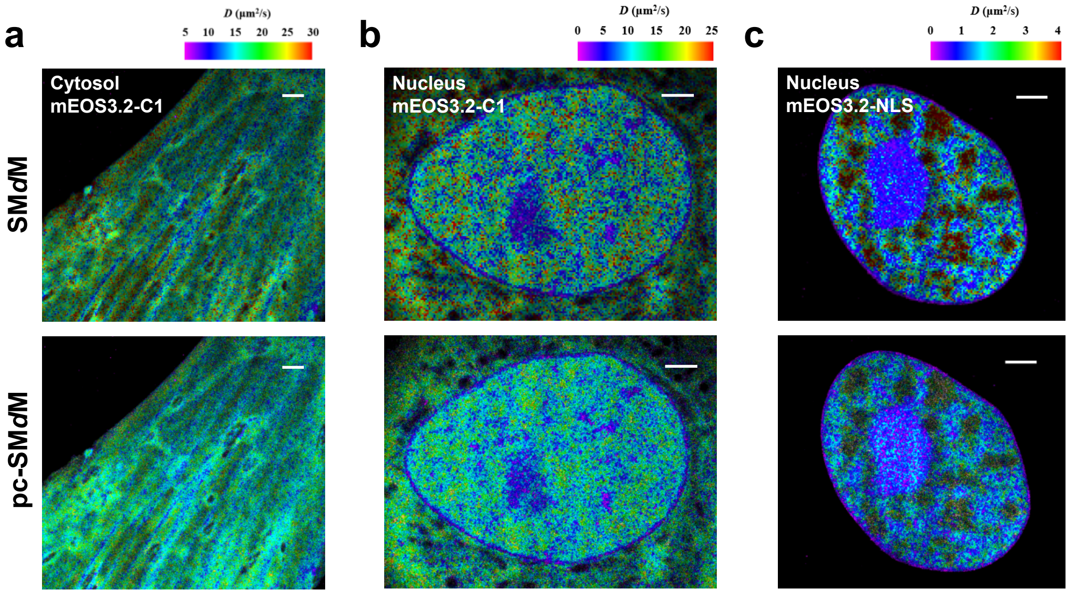


**Extended Data Fig. 9: Pixelated SM*d*M and pc-SM*d*M imaging of various mEos3.2 species with fast diffusion rates in live PtK2 cells.**

**a,** Pixelated SM*d*M and pc-SM*d*M images of mEos3.2-C1 in the cytoplasm. **b,** Pixelated SM*d*M and pc-SM*d*M images of mEos3.2-C1 in the nucleus. **c,** Pixelated SM*d*M and pc-SM*d*M images of mEos3.2-NLS in the nucleus. Scale bar: 2 μm. These results demonstrate that pc-SM*d*M can effectively map fast diffusion (>10 $\mu m^2/s$) in live cells, offering finer spatial resolution and maintaining the point-cloud data format for enhanced diffusivity analysis.
